# Two real use cases of FAIR maturity indicators in the life sciences

**DOI:** 10.1101/739334

**Authors:** Serena Bonaretti, Egon Willighagen

## Abstract

Data sharing and reuse are crucial to enhance scientific progress and maximize return of investments in science. Although attitudes are increasingly favorable, data reuse remains difficult for lack of infrastructures, standards, and policies. The FAIR (findable, accessible, interoperable, reusable) principles aim to provide recommendations to increase data reuse. Because of the broad interpretation of the FAIR principles, maturity indicators are necessary to determine FAIRness of a dataset. In this work, we propose a reproducible computational workflow to assess data FAIRness in the life sciences. Our implementation follows principles and guidelines recommended by the maturity indicator authoring group and integrates concepts from the literature. In addition, we propose a FAIR balloon plot to summarize and compare dataset FAIRness. We evaluated our method on two real use cases where researchers looked for datasets to answer their scientific questions. We retrieved information from repositories (ArrayExpress and Gene Expression Omnibus), a registry of repositories (re3data.org), and a searchable resource (Google Dataset Search) via application program interface (API) wherever possible. With our analysis, we found that the two datasets met the majority of the criteria defined by the maturity indicators, and we showed areas where improvements can easily be reached. We suggest that use of standard schema for metadata and presence of specific attributes in registries of repositories could increase FAIRness of datasets.

## Introduction

Data sharing and data reuse are two complementary aspects of modern research. Researchers share their data for a sense of community, to demonstrate integrity of acquired data, and to enhance quality and reproducibility of research [1]. In addition, data sharing is supported by the emerging citation system for datasets, scientific journal requirements, and funding agencies that want to maximize their return on investments in science [2], [3]. At the same time, researchers are eager to reuse available data to integrate information that answer interdisciplinary research questions and to optimize use of funding [4]. Although attitudes towards data sharing and reuse are increasingly favorable [1], data discovery and reuse remain difficult in practice [5]. Studies show that 40% of qualitative datasets were never downloaded, and about 25% of data is used only up to 10 times [6]. In addition, Vines et al. demonstrated that data availability decreases 17% per year due to the lack of appropriate hardware to access old storage media or because data were lost [7]. To be effective, data sharing and reuse need appropriate infrastructure, standards, and policies [5].

In 2016, the FORCE 11 group proposed guidelines to increase data reuse in the life sciences. These guidelines aimed to make data findable, accessible, interoperable, and reusable, and were summarized with the acronym FAIR [8]. In a short time the FAIR guidelines have gained remarkable popularity, and they are currently supported by funding agencies and political entities such as the European Commission, the National Institutes of Health in the United States, and institutions in Africa and Australia [9]. In addition, academic and institutional initiatives were launched to promote and implement data FAIRness, such as GOFAIR and FAIRsharing. Although largely adopted, the FAIR principles do not specify any technical requirement as they are deliberately intended as aspirational [9]. The lack of practical specifications generated a large spectrum of interpretations and concerns and raised the need to define measurements of data FAIRness. Some of the authors of the seminal paper proposed a set of FAIR metrics [10], subsequently reformulated as FAIR maturity indicators [11]. At the same time, they invited consortia and communities to suggest and create alternative evaluators. The majority of the proposed tools are online questionnaires that researchers and repository curators can manually fill to assess the FAIRness of their data (Table 1). However, the FAIR metrics guidelines emphasize the importance of creating “objective, quantitative, [and] machine-interpretable” evaluators [10]. Following these criteria, two platforms have recently been developed to automatically compute FAIR maturity indicators: FAIR Evaluation Services and FAIRshake. The first platform offers an evaluation of maturity indicators and compliance tests [11], whereas the second platform provides metrics, rubrics and evaluators for registered digital resources [12]. Both platforms provide use cases for FAIRness assessment, however they do not provide systematic analysis of evaluated datasets and repositories. Literature reports two studies evaluating FAIRness for large datasets. Dunning et al. [13] used a qualitative approach to investigate 37 repositories and databases. They assessed FAIRness using a traffic-light rating system that ranges from no to full compliance. Differently, Weber et al. [14] implemented a computational workflow to analyze the retrieval of more than a million images from five repositories. They proposed metrics specific for images, including time and place of acquisition to assess image provenance. The first study provides valuable concrete guidelines to assess data FAIRness, however the implementation was manual, differently from what the guidelines suggest. On the other side, the second study is a relevant example of computational implementation, although limited to retrieval of images and evaluation of 10 out of 15 criteria, and without unique correspondence between FAIR principles and maturity indicators.

**Table 1:** Online FAIR evaluators and studies in the literature assessing FAIRness of data repositories (the symbol ✓ indicates “yes”, the symbol – indicates “no”).

| Authors | Questionnaire / Platform | Manual Assessment | Automatic Assessment |  |  | Data / Code Repository |
| --- | --- | --- | --- | --- | --- | --- |
|  |  |  | Code / Language | Metadata Format | Protocol / Library |  |
| FAIRness evaluators |  |  |  |  |  |  |
| Wilkinsons et al. [10] | - | ✓ | - | - | - | GitHub |
| Australian Research Data Commons | FAIR self-assessment tool | ✓ | - | - | - | - |
| Commonwealth Scientific and Industrial Research Organization | 5 star data rating tool | ✓ | - | - | - | - |
| Data Archiving and Networked Services | FAIR enough? and FAIR data assessment tool | ✓ | - | - | - | - |
| GOFAIR consortium | FAIR ImplementationMatrix | ✓ | - | - | - | Open Science Framework |
| EUDAT2020 | How FAIR are your data? | ✓ | - | - | - | Zenodo |
| Wilkinsons et al. [11] | FAIR evaluation services | - | Ruby on Rails | JSON, Microformat, JSSON-LD, RDFa | nanopublications | GitHub |
| Clark et al. [12] | FAIRshake | - | Django and python | RDF | Extruct | GitHub |
| Studies assessing FAIRness of repositories |  |  |  |  |  |  |
| Dunning et al. [13] | - | ✓ | - | - | - | Institutional repository |
| Weber et al. [14] | - | - | python | DataCite | OAI-PMH | GitLab |
| Our approach | - | ✓ (partially) | Jupyter notebook with python | XML, JSON | request | GitHub |

In this paper, we propose a computational approach to calculate FAIR maturity indicators in the life sciences. We followed the recommendations provided by the Maturity Indicator Authoring Group (MIAG) [11] and we created a visualization tool to summarize and compare FAIR maturity indicators across various datasets and/or repositories. We tested our approach on two real use cases where researchers retrieved data from scientific repositories to answer their research questions. Finally, we made our work open and reproducible by implementing our computations in a Jupyter notebook using python.

## Materials and methods

### Use cases in the life sciences

We asked two available researchers in our department for a case where they looked for datasets in a scientific repository to answer a research questions. For each use case, the name used throughout the paper, research question, and investigated repository are:

- *Parkinsons_AE*: What are the differentially expressed genes between normal subjects and subjects with Parkinson’s diseases in the brain frontal lobe? To answer this question, the researcher looked for a dataset in the search engine of ArrayExpress, a repository for microarray gene expression data based at the European Bioinformatics Institute (EBI), United Kingdom [15];
- *NBIA_GEO*: What is the effect of the *WDR45* gene mutation in the brain? In this case, the researcher looked for a dataset in the search engine of Gene Expression Onmibus (GEO), a repository containing gene expression and other functional genomics data hosted at the National Center for Biotechnology Information (NCBI), United States [16].

### What is *data* and what is *metadata*?

The FAIR principles use the terminology *data*, *metadata*, and *(meta)data* (principles fully listed in Table 2). For our computational implementation, we needed precise definitions of these terms:

- *data*: According to the Merrian-Webster online dictionary, *data* are “information in digital form that can be transmitted or processed” [17];
- *metadata*: In the Merrian-Webster online dictionary, *metadata* are defined as “data that provide information about other data” [18];
- *(meta)data*: We interpreted it as *data and/or metadata*. We used *(meta)data* as:

- *data* for the principles R1, R1.1, and R1.2;
- *metadata* for the principles I1 and I3;
- *data and metadata* for the principles F1, F4, and A1.

**Table 2:**
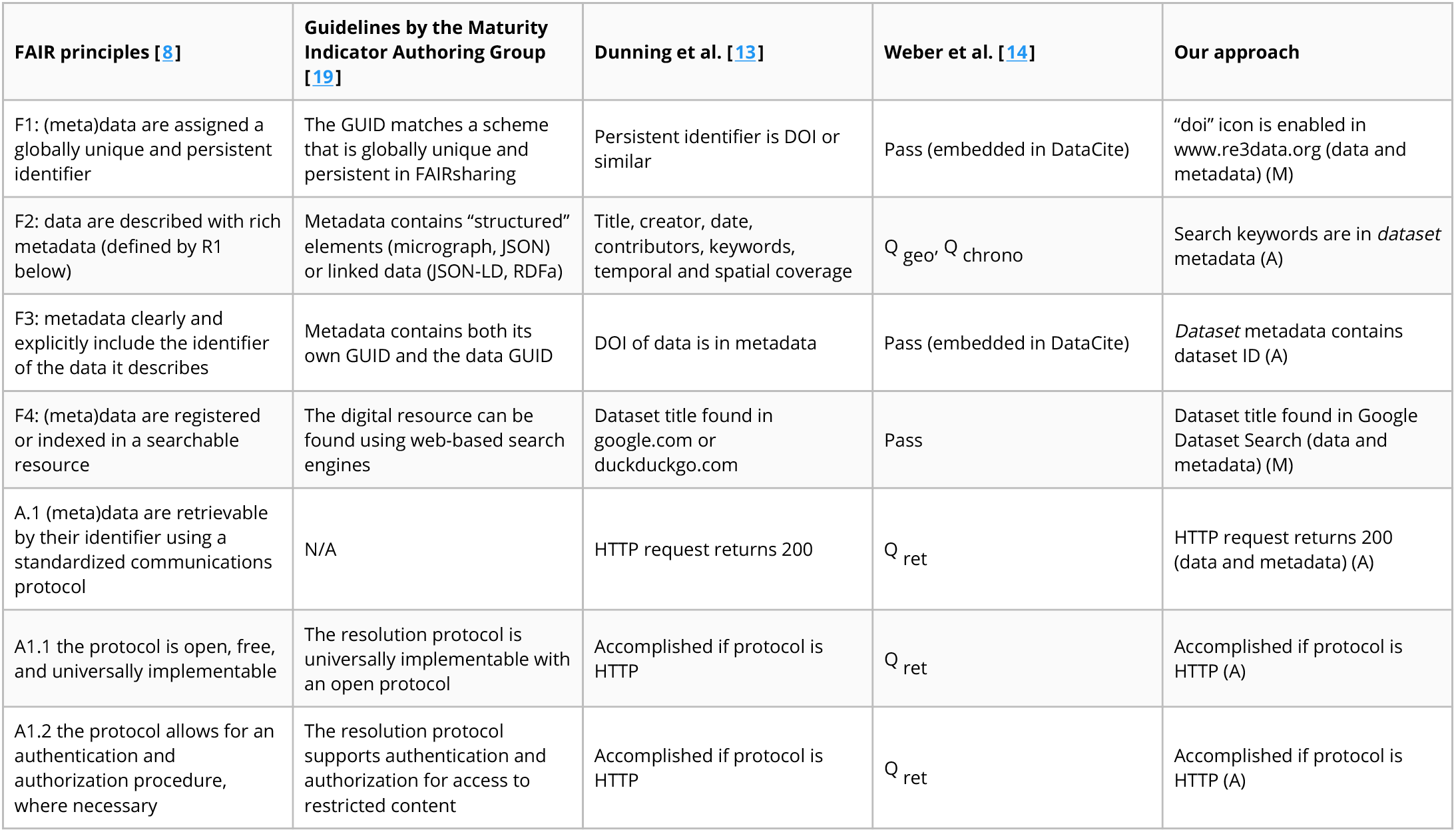
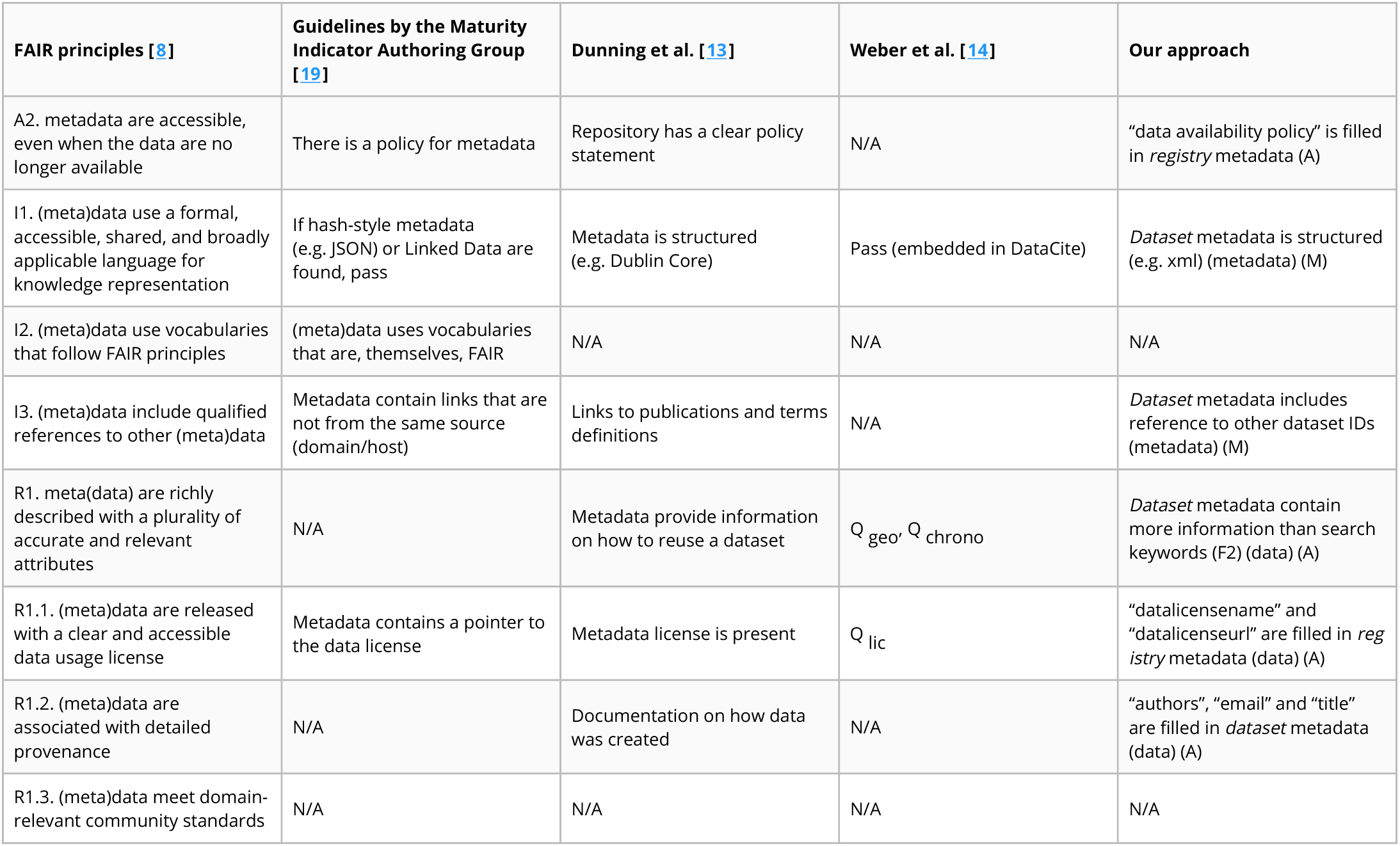
FAIR principles and corresponding evaluation criteria proposed by the Maturity Indicator Authoring Group [19], Dunning et al. [13], Weber et al. [14], and our approach. The criteria used in the first two works are extracted from their publication text, whereas the criteria by Weber et al. are from Table IV of their paper. The metrics Weber et al. developed are Q_geo_ for image location, Q_time_ for the time of picture acquisition, Q_ret_ when data is automatically downloadable only given its metadata, and Q_lic_ for found license. In our approach, *dataset* metadata refers to metadata retrieved from ArrayExpress and Gene Expression Omnibus, whereas *registry* metadata consists of metadata retrieved from re3data.org. In addition, we specify use of *(meta)data* as (data), (metadata), or (data and metadata), and automatic (A) or manual (M) procedure to retrieve information. Acronyms: GUID = Globally Unique IDentifier, DOI = Digital Object Identifier.

In our implementation, these terms assumed the following meaning:

- *data*: It is the actual dataset that researchers analyzed to answer their research question. The analysis of the dataset itself is out of the scope of this study;
- *metadata*: For the following principles, the corresponding *metadata* are:

- F2: Information that allow researchers to find the dataset s/he looks for. It coincides with the keywords used in the search;
- F3: Identifier of the dataset in the repository;
- I3: Reference to other metadata;
- R1: Information about the dataset, other than the search keywords;
- R1.1: Data license;
- R1.2: Data provenance as publication title, author names, and one author’s email address.

In all cases, we assumed that *data* and *metadata* were hosted in the same repository.

### Calculating FAIR maturity indicators

Because the FAIR guidelines emphasize on the importance of *data* and *metadata* being “machine-interpretable”, we collected information about datasets and repositories via an application programming interface (API) wherever possible. We queried three different sources:

- Data repositories (ArrayExpress and Gene Expression Omnibus): We programmatically queried each repository using the same keywords researchers had used in their manual query when looking for a dataset. From the obtained metadata, we retrieved information to calculate maturity indicators for the principles F2, F3, I1, I3, R1, and R12;
- Registry of repository: We queried re3data.org, a registry containing information about more than 2000 data repositories from various disciplines. We used the retrieved information to compute the maturity indicators for the principles F1, A2, and R12;
- Searchable resource: We queried Google Dataset Search, an emerging search engine specific for datasets, to quantify the principle F4.

The output of queries consisted of information structured in `xml`. Details about the computation of each specific maturity indicator are in Table 2 and in our Jupyter notebook (interactive on binder). To each maturity indicator, we assigned binary value 1 if the criterion was satisfied and 0 in the opposite case. The only exception was the maturity indicator F2, calculated as the ratio between the number of keywords in the metadata over the total number of keywords used by the researcher in the manual query, and thus ranging from 0 to 1. Similar to previous studies [19],[13],[14], we did not evaluate maturity indicators for the principles I2 and R1.3.

### Visualizing FAIR maturity indicators

To summarize and compare FAIRness of datasets, we developed the FAIR balloon plot using the R package ggplot2 [20]. In the graph, each row corresponds to a use case and each column to a FAIR maturity indicator. The size of each shape is the value of a specific FAIR maturity indicator for a particular dataset. Diamonds represent maturity indicators determined manually, circles depict maturity indicators established automatically, and crosses illustrate the maturity indicators we did not compute. Finally, colors represent the group of principles in the acronym: blue for findable, red for accessible, green for interoperable, and orange for reusable.

## Results

For both use cases, metadata contained all keywords used in the manual search (F2), dataset unique identifiers (F3), and additional information for data reuse (R1). In addition, they were structured in `xml` format (I1) and were released with a clear usage license (R11). The protocol used to retrieve all information was HTTP, which is standardized (A1), open, free and universally implementable (A11), and allows for authentication where needed (A12). In both cases, metadata were not assigned a persistent identifier (F1) and did not reference to other metadata (I3). Finally, the dataset of the use case *Parkinson_AE* was listed in Google Dataset Search (F4) and had detailed provenance (R12), whereas the dataset *NBIA_GEO* did not. Comparative summary of results is in Figure 1, and details of findings are in Table 3.

**Figure 1:**
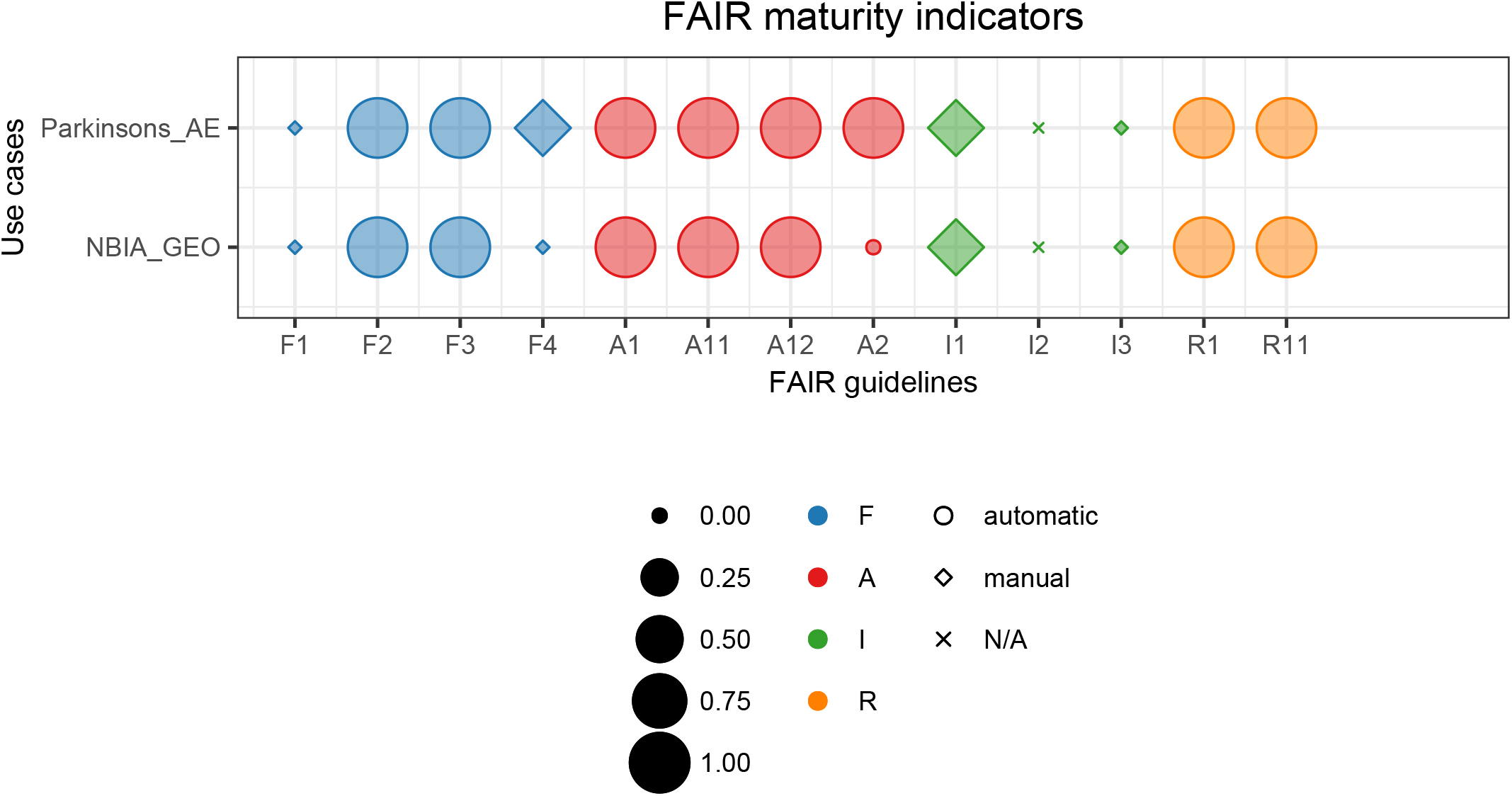
FAIR ballon plot. Comparative summary of FAIR maturity indicators for the two use cases evaluated in this work. Size corresponds to the numerical value of mutual indicators, colors represent FAIR categories, and shapes illustrate the way we retrieved information (N/A = not available). The graph can be fully reproduced from our Jupyter notebook on GitHub and interactively in binder.

**Table 3:** Comparison of API systems and FAIR maturity indicators for the two uses cases analyzed in this work. For each maturity indicator, we indicate the outcome in natural language and in numbers (1 for pass and 0 for fail).

| Use case | Parkinsons_AE | NBIA_GEO |
| --- | --- | --- |
| Repository / Database | Array Express | Gene Expression Omnibus |
| Search output on browser | link | link |
| API |  |  |
| Type | REST | REST |
| Documentation | link | link |
| Output format | XML | XML |
| FAIR maturity indicators |  |  |
| F1 (Persistent identifier) | No (0) | No (0) |
| F2 (Findable metadata) | parkinson's disease, normal, homo sapiens, transcription profiling by array, raw data, frontal lobe, male, female (1) | nbia, homo sapiens, expression profiling by array (1) |
| F3 (Unique identifier) | 219251 (1) | 200070433 (1) |
| F4 (Google Dataset Search) | Yes (1) | No (0) |
| A1 (Communication protocol) | request status code = 200 (1) | request status code = 200 (1) |
| A11 (Open and free protocol) | Yes (1) | Yes (1) |
| A12 (Communication protocol) | Yes (1) | Yes (1) |
| A2 (Metadata always accessible) | Yes: https://www.ebi.ac.uk/arrayexpress/help/data_availability.html (1) | No (0) |
| I1 (Language representation) | XML (1) | XML (1) |
| I2 (FAIR vocabularies) | Not evaluated (None) | Not evaluated (None) |
| I3 (Reference to other metadata) | No (0) | No (0) |
| R1 (Metadata for reuse) | 56 metadata fields (1) | 58 metadata fields (1) |
| R1.1 (License) | name: other<br>url: https://www.ebi.ac.uk/arrayexpress/help/data_availability.html (1) | name: other<br>url: http://www.ncbi.nlm.nih.gov/geo/info/disclaimer.html (1) |
| R1.2 (Provenance) | Authors: Garcia-Esparcia P, Schlüter A, Carmona M, Moreno J, Ansoleaga B, Torrejón-Escribano B, Gustincich S, Pujol A, Ferrer I<br>Title: Functional genomics reveals dysregulation of cortical olfactory receptors in parkinson disease: novel putative chemoreceptors in the human brain (1) | No (0) |
| R1.3 (Community standards) | Not evaluated (None) | Not evaluated (None) |

## Discussion

We proposed a semiautomatic computational workflow to evaluate FAIR maturity indicators for scientific data repositories in the life sciences. We tested our method on two real use cases where researchers looked for datasets to answer their scientific questions. The two cases scored similarly. Finally, we created a FAIR balloon plot to summarize and compare our results, and we made our workflow open and reproducible.

Real use cases in the life sciences were the starting point of our computational implementation. In their guidelines, the MIAG suggests to calculate maturity indicators starting from a global unique identifier (GUID) (e.g. InChI, DOI, Handle, URL) [19]. However, a priori knowledge of a GUID often signifies that a researcher has already found and accessed the dataset s/he is going to reuse. In addition, it assumes that the repository of interest provides unique identifiers, which is not the case for ArrayExpress and Gene Expression Omnibus, based on the information we retrieved from re3data.org. Similar to Weber et al. [14], we decided to start our computations from dataset retrieval. We asked two researchers in our departments to show us how they looked for the datasets of interest and which keywords they used. Then, we computationally reproduced their manual search by programmatically retrieving data and metadata using their same keywords. We recognize that this approach limits the generalization of FAIRness calculation. While creating a use case for every dataset is extremely demanding, the same dataset could be used to answer different research questions. However, we think that an exhaustive set of real use cases could provide valuable insights on how to practically achieve data FAIRness, as demonstrated in the literature [21].

To assess data FAIRness, we implemented criteria that follow principles and guidelines recommended by the MIAG [19], reused concepts from similar studies in the literature [13],[14], and added new considerations (Table 2):

- *Findability*: The criteria to assess principles F1 (unique identifier), F3 (metadata includes identifier), and F4 ((meta)data are indexed) are similar for all authors. In our case, to assess F1 we investigated whether a repository provides DOI in the registry re3data.org. We chose this registry because it is one of the largest registries of scientific repositories, and it provides an open API. For F3, we accepted any dataset identifier provided by the repository as the principle does not explicitly mention restrictions on the characteristics of the identifier. Finally, for F4 we looked for dataset titles in Google Dataset Search. We chose this searchable resource because it could become one of the main search engines specific for data in the future, similar to Google Scholar for publications. Opposite to the previous maturity indicators, the implementation of F2 (data are described with rich metadata) has large variations across authors. The MIAG recommends to evaluate whether metadata contains “structured” elements, Dunning et al. looked for attributes that favor findability, whereas Weber et al. used metrics of time and space of image acquisition. We followed the criteria suggested by Dunning et al. and looked for the keywords that researchers had used in their manual search to *find* datasets.
- *Accessibility*: Similar to the other authors, we retrieved our data using the HTTP protocol, which is free, open and allows for authentication, and thus satisfies all the requirements of the A1 group. Also, there is concordance among authors for the principle A2, which requires that a repository should explicitly provide a policy for data availability. In our implementation, we looked for the policy in re3data.org.
- *Interoperable*: Similarly to the MIAG, we assigned a positive score to metadata in a structured file format, such as `xml` (I1). On the other side, Dunning et al. and Weber et al. suggested that metadata should be in a standardized schema, such as Dublin Core or DataCite, which would increase data interoperability and simplify retrieval. None of the studies assessed I2 (vocabularies are FAIR) because it would require a separate implementation that includes the recursive nature of the FAIR principles. Finally, for I3 all authors looked for references to other datasets in metadata.
- *Reusable*: Although the MIAG does not provide any guideline, authors implemented different ways to assess R1 (plurality of relevant attributes). While Weber et al. used the same metrics for F2, Dunning et al. focused on metadata that provide information on how to reuse a dataset. In our implementation, we assess the presence of metadata attributes other than search keywords. The principles R11 (availability of data usage license) and R12 (data provenance) had a straight-forward implementation for all authors. In our approach, we looked for a data license in re3data.org and for authors, author emails, and title of the corresponding publication in the metadata from the dataset repository. Finally, none of the authors evaluated whether metadata follow community standards (R13), as community agreements are not formally established yet.

We assessed FAIR maturity indicators using a mixed manual and automatic approach. In the literature, Dunning at al. used a fully manual approach to assess the maturity indicators, whereas Weber et al. used a completely automatic approach, calculating 10 of the total 15 maturity indicators. Our mixed approach enabled automatic assessment of maturity indicators wherever possible, and to manually complement when we could not retrieve information via API. Because repositories do not use a standardized metadata schema, our mixed implementation required prior manual investigation of metadata attributes for each repository. For example, ArrayExpress uses the attributes “authors”, “email”, and “title” that we could use for the principle R12, whereas Gene Express Omnibus does not have attributes for provenance.

To summarize and compare dataset FAIRness, we created a FAIR balloon plot. As the MIAG guidelines recommend, we did not create a final score to avoid concerns for data and resource providers [11]. In our plot, we combined colors, sizes, and shapes of graphical elements to provide a summary of principles, scores, and type of information retrieval (manual, automatic, not assessed) for each dataset. In this visualization, a dataset that reached full FAIRness would have all maturity indicators depicted as circles with maximum size, meaning full score and automatic retrieval. In addition, by vertically stacking representations for different datasets, we can visually compare FAIRness levels for each maturity indicator. In the literature, another example of visualization are *insigna*, created for the platform FAIRshake [12]. They consist of multiple squares colored from blue (satisfactory) to red (unsatisfactory) for different levels of FAIRness. In addition, they can dynamically expand to visualize multiple scores calculated using different rubrics (i.e. criteria). Although this representation embeds the possibility of using different criteria, it does not allow direct comparison across datasets. Finally, we applied our FAIR balloon plot to the results collected by Dunning et al. to demonstrate that this kind of visualization can be reused for FAIR assessment with other criteria (Figure 2).

**Figure 2:**
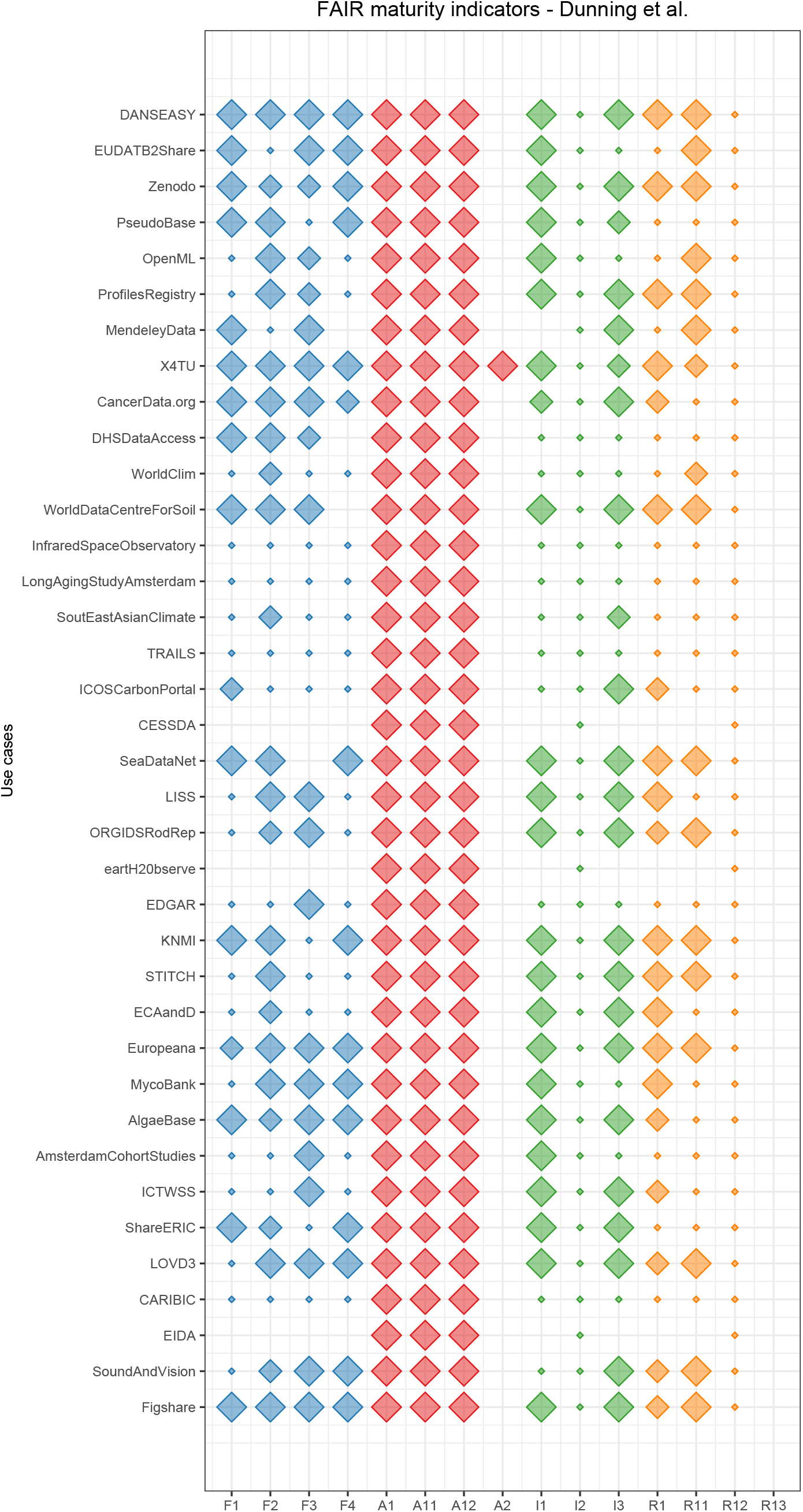
FAIR balloon plot for the repositories analyzed by Dunning et al. [13] (data available at their institutional repository). From their quantitative scores, we converted “complies completely” to 1, “just about/maybe not” to 0.5, and “fails to comply” to 0. We did not assign any value to “unclear”, which is thus represented as missing elements. The graph can be fully reproduced from our Jupyter notebook on GitHub and interactively in binder.

To make our analysis open and reproducible, we implemented our workflow in a Jupyter notebook. However, changes to APIs or metadata attributes could affect reproducibility of the results. The possibility of querying a specific version of a repository could be a possible solution. In addition, we implemented our approach in python, a language increasingly used in various scientific communities that can potentially favor extension and reuse of our work. For new datasets, FAIR maturity indicators could be evaluated by changing the search procedure and the values assigned manually.

The two analyzed datasets (*Parkinsons_AE* and *NBIA_GEO*) met the majority of the criteria used to assess FAIRness. Higher FAIRness compliance could be reached by using a standard schema for metadata (e.g. Dublin Core, DataCite, or schema.org), which could include all attributes required by the principles, and by providing explicit information about data policy, licenses, etc. to registries of repositories.

In conclusion, we proposed a reproducible computational workflow to assess data FAIRness in the life sciences, and we created a FAIR balloon plot to summarize and compare FAIRness compliance. We evaluated our approach on two real cases, and we demonstrated that the FAIR balloon plot can be extended to other FAIRness analyses. Finally, we suggested that use of standard schema for metadata and presence of specific attributes in registries of repositories could increase FAIRness of datasets.

## Acknowledgments

This work received funding from from the European Union’s Horizon 2020 research and innovation programme via NanoSolveIT Project under grant agreement No 814572 and via RiskGONE Project under grant agreement No 814425. We would like to thank Nasim B. Sangani (0000-0002-3560-2058), Gwen Keulen, and Friederike Ehrhart (0000-0002-7770-620X) for the use cases, Tobias Weber (0000-0003-1815-7041) for the insightful discussion about data retrieval, and Lauren Dupuis (0000-0003-2606-3045) for revising our manuscript.

We created this manuscript using manubot [22].

